# The architecture of multifunctional ecological networks

**DOI:** 10.1101/2023.07.02.547400

**Authors:** Sandra Hervías-Parejo, Mar Cuevas-Blanco, Lucas Lacasa, Anna Traveset, Isabel Donoso, Ruben Heleno, Manuel Nogales, Susana Rodríguez-Echeverría, Carlos Melián, Victor M. Eguíluz

**Author notes:** SHP and MCB are both first authors. LL and VME are both corresponding authors.

## Abstract

Understanding how biotic interactions affect ecosystem functioning has been a research priority in natural sciences due to their critical role in bolstering ecological resilience^1–3^. Yet, traditional assessment of ecological complexity typically focus on species-species effective interactions that mediate a particular function (e.g. pollination^4^ or seed dispersal^5^), overlooking the synergistic effect of multiple functions that further underpin species-function and function-function interactions in multifunctional ecosystems. At the same time, while ecological network theory holds a potential to quantify the relationship between biodiversity and ecosystem multifunctionality^6, 7^, its connection has been done mainly conceptually, due to challenges measuring different interactions and establishing their relevance across multiple niche dimensions^8, 9^. Such lack of quantitative studies therefore limits our ability to determine which species and interactions are important to maintain the multiple functions of ecosystems^10^. Here we develop a framework –derived from a resource-consumer-function tensor analysis-that bridges these gaps by framing biodiversity-ecosystem multifunctionality in terms of multilayer ecological network theory. Its application to recently collected ecological data –– reporting weighted interactions between plants, animals and fungi across multiple function types––allows to (i) unveil and quantify the existence of both (multi-functional) keystone species and a dual function keystoneness pattern, and (ii) project plants and functions into a similarity space where clear clusters emerge and the importance of weak links is manifested. This dual insight from species and functional perspectives will better guide conservation efforts to reduce biodiversity loss.

## Main

All species are permanently involved in a myriad of entangled interactions with other coexisting species^11–13^, playing multiple and simultaneous ecological roles that together define the multiple dimensions of their Eltonian niche^14, 15^. In a nutshell, ecosystems are inherently multidimensional complex systems, and different types of collective effects throughout the complexity hierarchy^16^ are therefore expected to emerge, including not only effective species-species interactions but also^11, 12^: species-function interactions (e.g. a herbivore interfering pollination, plant species participating in various functions in heterogenerous ways) and effective function-function (e.g. an animal pollinating and dispersing the same plant species) ones. Thus, any representation aiming to capture the relationship between biodiversity and ecosystem multifunctionality^6, 7^ requires incorporating such complexity in a meaningful way, i.e. it needs to enrich biodiversity-ecosystem multifunctionality^10^ patterns by accounting for species-rich interactions and functions^17, 18^. From a modelling perspective, while traditional mathematical frameworks such as network theory have proven useful to gain insights on, for instance, extinction cascades^19, 20^, there is a need to embrace more sophisticated ones –e.g. multilayer networks^21–23^ in order to naturally incorporate different types of interactions^24^ (both direct and indirect) into a unique modelling framework. At the same time, accessing the necessary fine-grained empirical observations of species interactions is inherently a difficult task^25^, often resulting in incomplete observations or indirectly-inferred^26, 27^ data. In addition, multilayer modelling requires stringent standardised measurements of species’ importance in each function (i.e. using the same currency to quantify different types of interactions), thereby hampering advances in the field. Integrating several ecological functions, thus, represents both a necessity and a challenge^28^ given the difficulties imposed by data collection and standardisation, and their subsequent translation into appropriate modelling paradigms. As a result, there is a lack of quantitative studies connecting biodiversity-ecosystem multifunctionality to ecological networks^8, 29^.

Here we take the first steps to bridge this gap, both from a theoretical and experimental viewpoints. Our modelling approach is inspired by the consumer-resource paradigm^30, 31^, whereby plants are seen as “resources”, and “consumers” encapsulate different types of animals or fungi involved in both mutualistic and antagonistic interactions (note, however, that the methodology and analysis can be generally extended to other systems where a consumer-resource paradigm exists, such as economic systems^32^). By extending multiple consumer-resource interactions to many functions, the biodiversity-ecosystem multifunctionality relationship can therefore be analysed not only from a multiple interaction but also from a multiple function perspective. Then, by leveraging the relative simplicity of a small island ecosystem –Na Redona, in the Balearic Islands (W Mediterranean Sea)-, we apply such a framework to unveil the multifunctional architecture of an ecological system reconstructed via fieldwork data encompassing a total of 1537 weighted interactions between plants, animals and fungi across six ecological functions (layers). Our common species in all six layers are plants, which interact with pollinators, herbivores, seed dispersers, and three types of (pathogenic, saprotrophic and symbiotic) fungi.

Operationally, we will first proceed to integrate the multiple dimensions of ecological interactions into a Resource-Consumer-Function tensor (hereafter RCF). The RCF can be visualised as a (multipartite) multilayer weighted network. The architecture of MultiFunctional Ecological Networks (hereafter MFEN) can be decoded from the RCF itself by suitably contracting the consumer index and yielding a resource-function matrix or map. We will show that the representation of the MFEN in the Na Redona system displays a nested pattern^33^. As MFEN studies the architecture of species-to-function interactions, interpreting such pattern necessarily leads to the existence of both (multifunctional) keystone species and, notably, also ‘function keystoneness’, a new concept we coin and discuss here. As a matter of fact, while all ecological functions are important, one can indeed ask whether their roles as ecosystem assemblers are similar or not, and how interactions and functions connect each other, to form a keystone interaction-function core containing the most critical functions for ecosystem functioning. We propose that this perspective aligns with approaching the ecosystem through a function-centric lens, rather than a traditional phytocentric one. Indeed, just as keystone species encode, among other properties, robustness (i.e. the ability to maintain performance in the face of perturbations) and resilience^34^, the response of the ecosystem to some disturbances may also occur in the functional dimension. For instance, non-native herbivores can disrupt chemically-mediated interactions between plants and herbivores, pollinators, predators, and parasites that respond to herbivore-induced plant volatile cues^35^. Because there is a wide variety of interactions involving native organisms that a single non-native could potentially impact, the challenge lies in determining which functions should be prioritised. Therefore, defining and identifying key ecological *functional cores* –critical for proper ecosystem development and balance– and studying the robustness and adaptation of these functional cores to disturbances can be crucial to understanding ecosystem functioning and resilience^36^.

Below, we illustrate how distinct methods applied to the original RCF tensor directly enable us to address and quantify a wide range of significant ecological questions within a unified methodological framework. These inquiries encompass determining multifunctional species keystoneness, the dual challenge of function keystoneness, and keystone functional cores to assess ecosystem clustering and robustness against perturbations of species or functions, and even establishing a resource-function mapping. Each of these questions indeed mirror different facets of the polyhedric structure of the ecosystem (see **Fig. 1** for an illustration of each step of the framework). Application of the proposed framework to data gathered in Na Redona islet unveils an intertwined keystoneness hierarchy both in the species and function dimensions that overall provide a better understanding of the complex structure of ecosystems.

**Figure 1.**
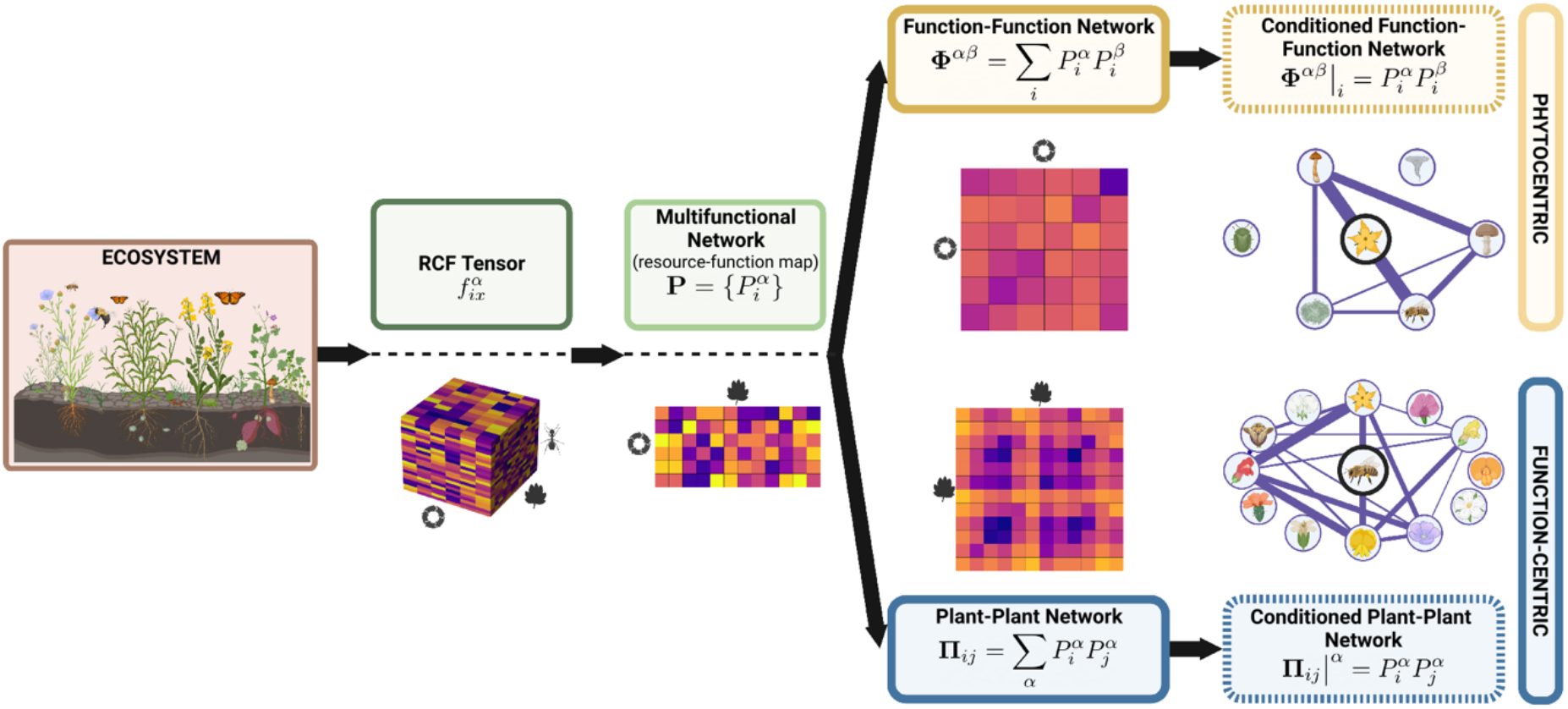
Conceptual framework. Ecosystems are multidimensional complex systems whose interactions are captured by a rank-3 Resource-Consumer-Function tensor (here, a plant-animal/fungi ecological interaction tensor). Contracting consumers out, we obtain a resource-function matrix or map –the MultiFunctional Ecological Network (MFEN)– that encapsulates how plant species and functions are intertwined in the ecosystem. Further projections of MFEN yield function-function networks that characterise how each plant species participate and intertwine different functions (phytocentric embedding) and plant-plant networks that characterise how each ecological function participates and intertwines different plant species (function-centric embedding).

### Complete data representation: the Resource-Consumer-Function tensor (RCF)

The precise span of dimensions is driven by the extent of our data (see Methods), which includes direct observation of 16 plant species, 629 fungal ASV (i.e. amplicon sequence variants) and 46 animal species, interacting across six fundamentally different ecological functions (pollination, herbivory, seed-dispersal, decomposition, nutrient uptake, fungal pathogenicity). The complete relational dataset is thus formalised in terms of a rank-3 tensor 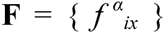 that we call the RCF. Interpreting the architecture of this tensor as a network^21^, we see that RCF displays two groups of nodes: resource nodes (plant species, denoted by Latin letters *i, j*), and consumer nodes (different animal and fungi species, denoted by the Latin letter *x*). By construction, such network is *multipartite*, since interactions (links) take place between groups, but no direct intragroup is directly recorded. Furthermore, links characterise different types of interactions, and, therefore, the network is naturally a *multilayer* one^22^, where each layer represents the wiring architecture according to a concrete function (functions are denoted by Greek letters α, β). Finally, such network is *weighted*: for each layer α, data allows us to calibrate the strength of the interaction between a resource node (plant species *i*) and a consumer node (animal/fungi species *x*), resulting in the link weight 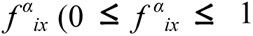, see Methods).

The RCF can be effectively visualised as a multipartite edge-coloured weighted network (see **Fig. 2**, after having processed the layout via the Infomap community detection algorithm; details provided in Methods), revealing that consumers are often centred around a single plant species, forming clusters (see, however, the cluster consisting of *Lavatera maritima* and *Geranium molle*). Interestingly, cross-cluster links are also present, depicting that the ecosystem is formed by many clusters connected via an animal/fungi consortium. For alternative visualisations of the RCF tensor, see Extended Data Figure **EDF1**.

**Figure 2.**
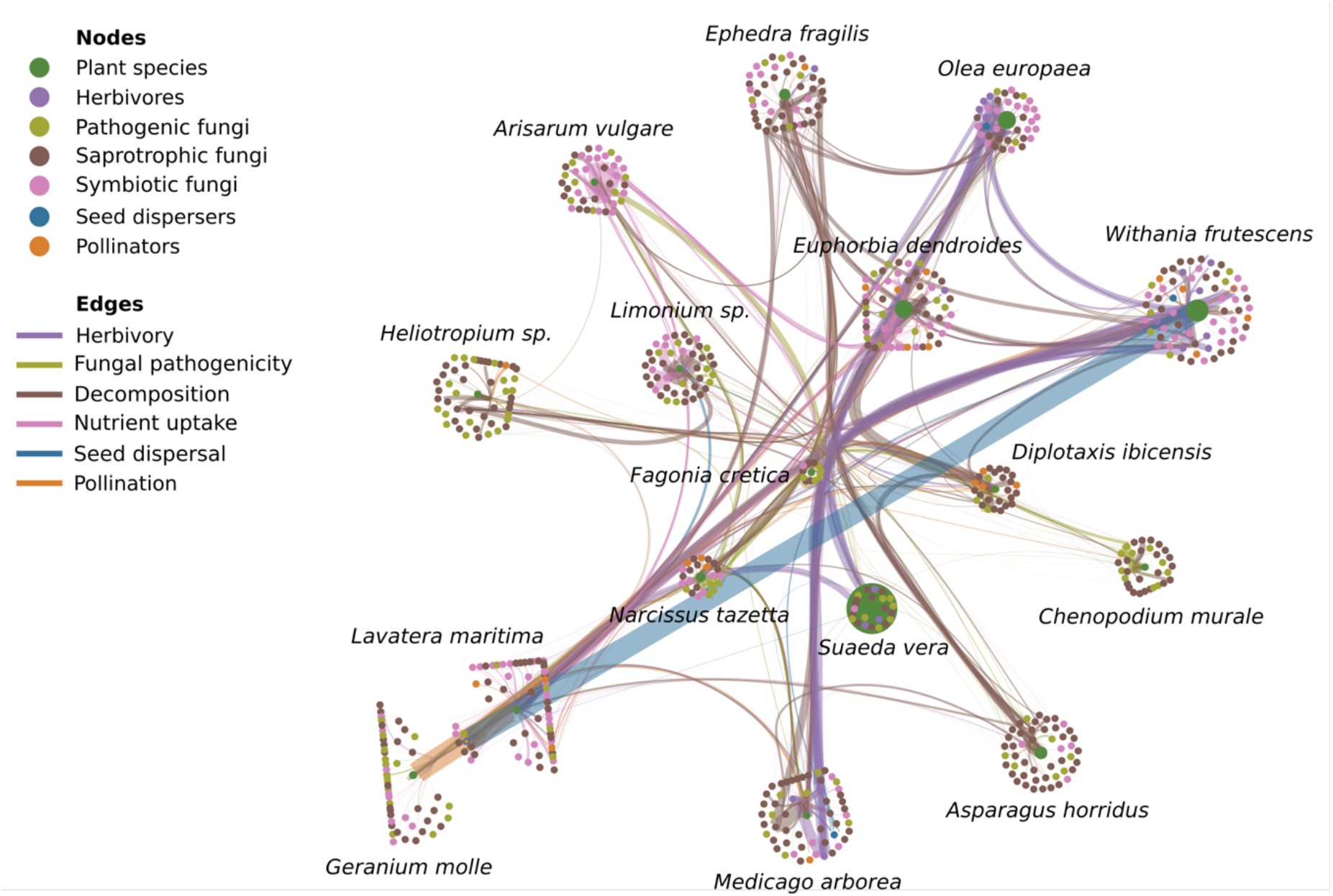
Basic visualisation of the Resource-Consumer-Function tensor (RCF) from the Na Redona dataset. There are 691 nodes (16 plant species and 675 animal/fungus species) interacting via six functions: pollination, herbivory, seed-dispersal, decomposition of plant matter, nutrient uptake (mycorrhizas), and fungal pathogenicity, for a total of 1573 weighted interaction links. Node colours account for plant species (green) and animal/fungus species (colour-coded according to the primary function they are involved in). The sizes of plant-nodes represent their observed abundance in the field (see Methods). Link widths quantify the weight of interaction and link colours denote the functional interaction type. In this suitable layout, species are clustered via Infomap’s community detection algorithm after flattening the RCF (for other layouts, see **EDF1**). Fifteen out of the sixteen clusters are comprise a single plant species, while one contains two plant species (Geranium molle and Lavatera maritima). Animals/fungi are associated with groups of plants to form a consortium to deliver the functionality of the ecosystem.

### The Multifunctional Ecological Network (MFEN)

To quantify the relationship between species and ecological functions the system embodies, we ‘contract’ the consumer indices into the RCF tensor, thereby building a *resource-function matrix,* which is the weighted adjacency matrix of the (bipartite) Multifunctional Ecological Network (MFEN). The weights of this bipartite network 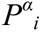 are called *participation strengths* and account for the probability that *i* participates in function α. To These are estimated from the empirical values of 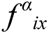, and making the parsimonious assumption of no correlation between consumers, which yields 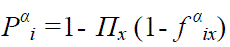 (see Methods for the derivation and **Fig. 3a** for an illustration). This choice for computing 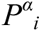 is parsimonious and can be refined to incorporate correlations (e.g. via the co-visitation patterns of two or more pollinator species) if empirical evidence of such correlation kernels is eventually available (e.g. via phylogenetic, trait similarity or field).

**Figure 3:**
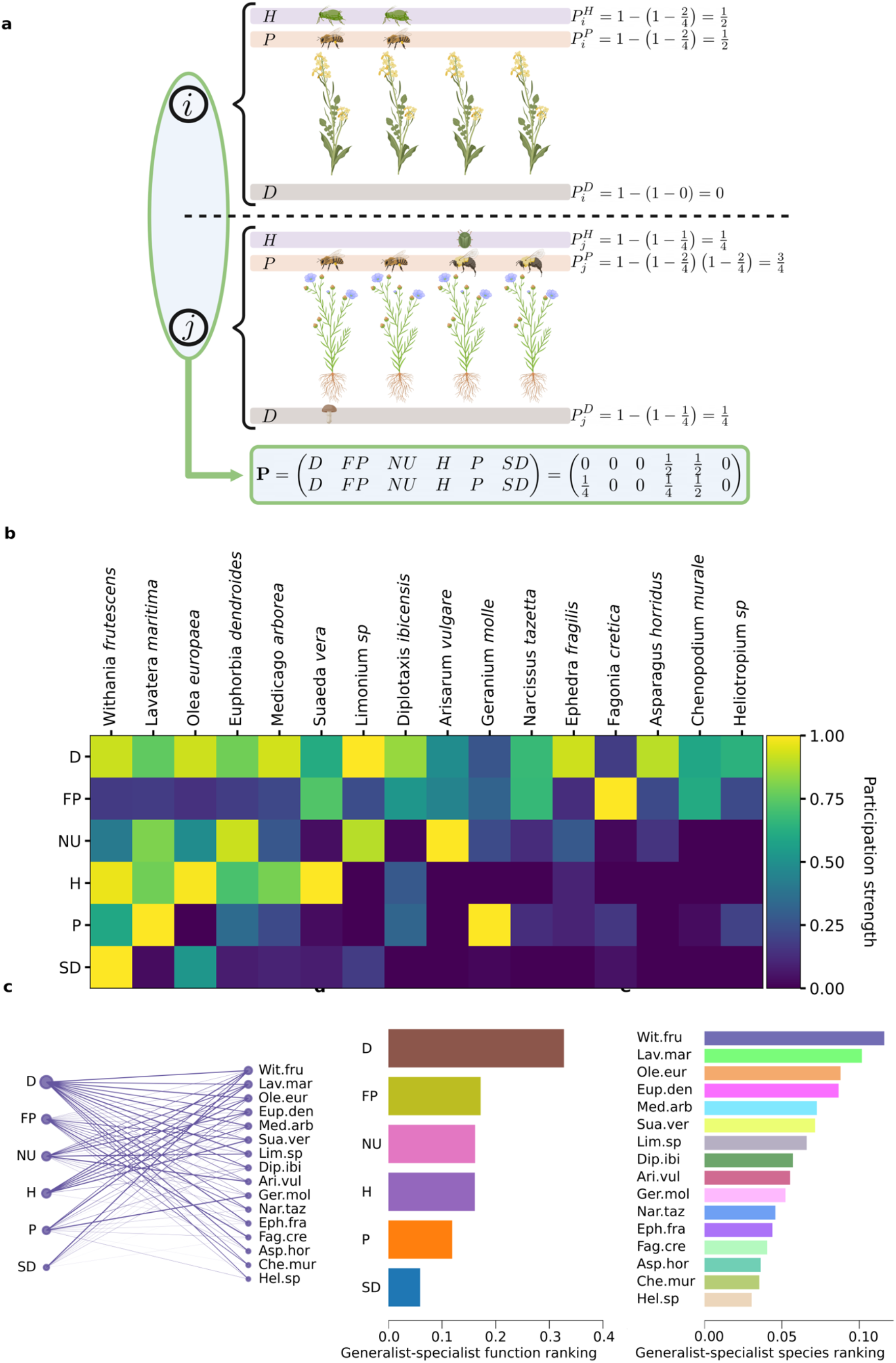
**Nestedness in The MultiFunctional Ecological Network of the Na Redona dataset**. (a) Illustrative cartoon of how the MultiFunctional Ecological Network **P** whose entries are the participation strengths P^α^_i_ is computed: for plant species i, two individuals have been e.g. pollinated and participated in herbivory function (see Methods). (b) Concrete matrix **P** obtained for the Na Redona dataset, showing a stylised nested structure, which suggests that (i) plant species rank in a generalist-specialist dimension (in terms of how many functions and hhow strongly they participate in) and that (ii) functions rank in a generalist-specialist dimension (in terms of how many species and how intensily are involved in a given function). (c) The MultiFunctional Ecological Network of the Na Redona dataset, visualised as a bipartite (resource-function) network. (d-e) Basic generalist-specialist function rankings of functions (d) and plant species (e). By evaluating the strength contribution of nodes within each function relative to the overall strength of all nodes, we were able to determine the relative importance of each species and function within the network (see Methods).

The MFEN is thus given by the matrix 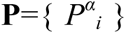, where *i* ranges across plant species and α ranges across functions^a^. The resulting **P** obtained with our empirical data from Na Redona is presented in **Fig. 3b** (a visualisation of the bipartite network is also provided in **Fig. 3c)**. **P** clearly displays (see below) a stylised nested structure, as commonly found in, for example, mutualistic interaction networks, world trade, inter-organisational relations, and others^33^, although in this case such pattern notably emerges in a species-to-function setting.

Observing such a nested pattern firstly indicates that the participation of plant species throughout different ecological functions is heterogeneous. To classify species as ecological function-generalists or function-specialists, it is crucial to consider both the number of functions in which they participate, and the strength of their participation^37^. For instance, *Withania frutescens* and *Lavatera maritima* participated in all six functions with an average probability of 0.6 or greater, indicating they act as generalist species. In contrast, others such as *Chenopodium murale* and *Heliotropium sp*. emerge as more “function-specialists”, participating in four out of the six functions with lower average probabilities (0.3 or less) (see **Fig. 3e** for a simple preliminary quantification). The emergence of such hierarchy suggests the existence of *multifunctional keystone plant species*, as we will fully develop and quantify below.

Second, the non-trivial nested pattern also suggests the existence of functions that are ‘plant–generalist’ –meaning they are participated by many plant species with systematic high participation strength-. This is for instance the case of decomposition –via saprotrophic fungi-, with an average participation of 0.7 by all plant species. This can then be compared to other functions, which are participated by fewer plant species with weaker strength, such as seed-dispersal –with an average participation of 0.2 by 9 of the 16 species– (**Fig. 3d** provides a preliminary quantification). Interestingly, finding a heterogeneous degree of participation opens up the question of whether and how robust an ecosystem is with respect to perturbations to functions (instead of species as commonly assumed), and overall naturally leads to formulating the novel concept of *function keystoneness*, that we will also fully develop and quantify later.

### Phytocentric embedding: function-function networks and Multifunctional plant species keystoneness

We now dive deeper into the multifunctional species keystoneness initially identified in **Fig. 3e**, and into the role that plant species play as ecosystem assemblers (i.e. the phytocentric perspective). To this aim, we proceed to project MFEN into the function class and thus extract a function-function effective interaction network with *N=6* nodes and weighted adjacency matrix 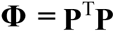 (superindex T denotes matrix transposition) that leverages how species connect –actually, broker– functions in the ecosystem. Mathematically, **Φ** allows for several equivalent interpretations. First, the elements 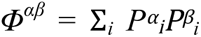 enumerate the effective paths (passing via a resource-species) connecting two functions α and β, and by weighting such paths, effectively compute the expected number of paths connecting α and β. This is therefore based on the number of plant species that simultaneously participate in both functions α and β, and hence quantifies the role of plant species as function assemblers. Second, interpreting the different participation strengths as function features, **Φ** adopts the mathematical form of a correlation matrix, i.e. a similarity-based matrix (see Methods), and quantifies how similar functions are to each other based on their participation strength pattern across plant species, i.e., in a plant-feature embedding. See **Fig. 4a** for a visualisation of **Φ** of the Na Redona dataset, showing that edge weights are indeed heterogeneous, indicating that plants impact functions assembly/homophily in a non-trivial way.

**Figure 4:**
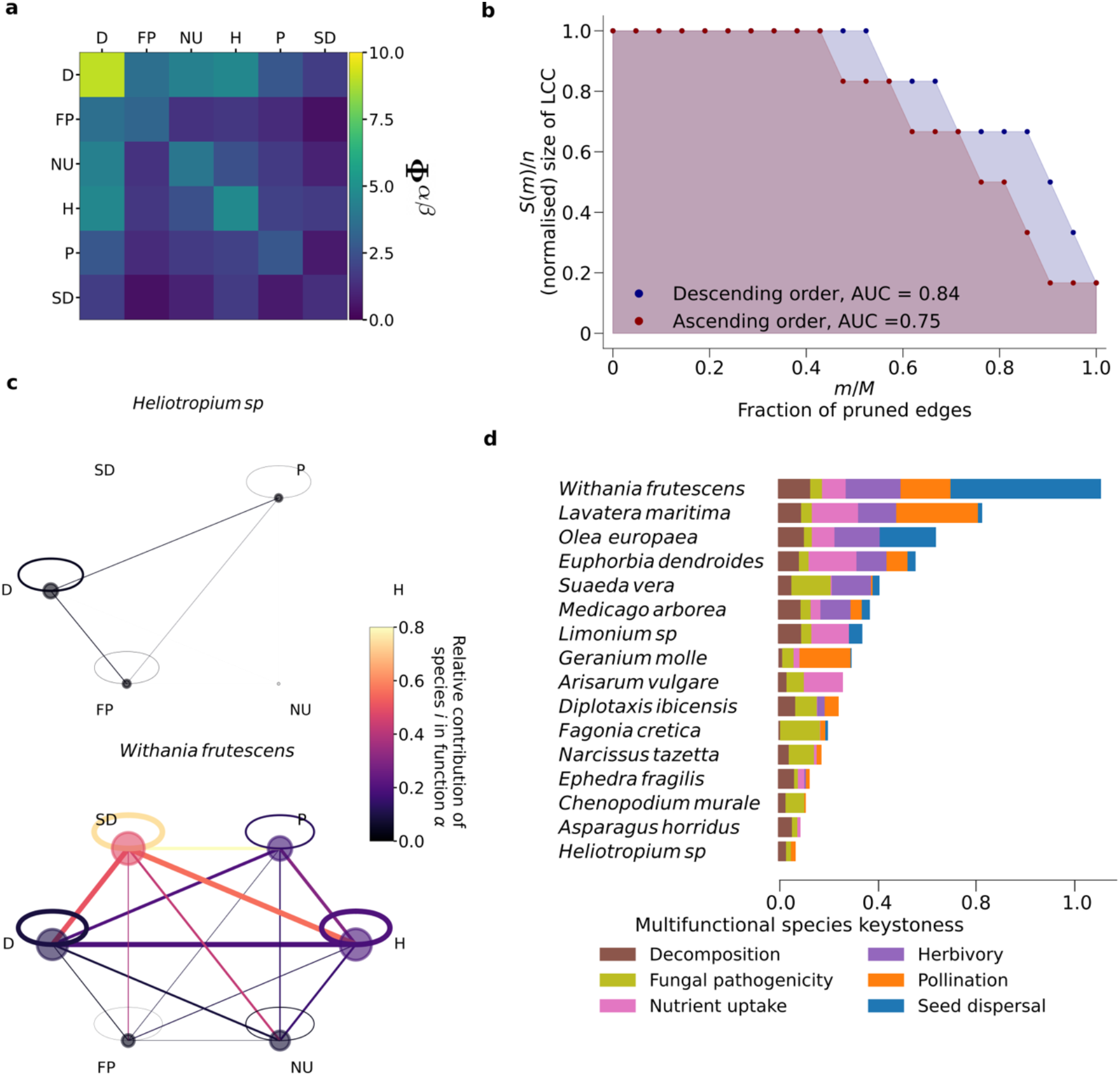
Phytocentric embedding: function-function networks and multifunctional species keystoneness. (a) Function-function network’s weighted adjacency matrix **Φ** for the Na Redona dataset. (b) Pruning analysis of **Φ**, plotting S(m)/n, the (normalised) size of the largest connected component of the pruned **Φ** as a function of the fraction of deleted edges m/M, for two pruning strategies (removing edges in decreasing (blue) or increasing (red) order of link weights, respectively). The Area Under the Curve (AUC) is a measure of the network’s robustness to pruning, showing that it is overall robust (large AUC values), but more sensible to deletion of links with lower weight (weaker ties) as this pruning strategy results in a lower AUC. (c) Two examples of conditioned function-function networks **Φ**|_’_ computed by conditioning on the plant species Withania frutescens (bottom) and Heliotropium sp (top), showing two cases of plant species with very different multifunctional roles in the ecosystem. Node colours represents each species’ relative contribution to the total strength of each function. For a species i, we obtain its contribution to function α relative to that of all species by summing the edge weights (strength) of node i in conditioned function-function network **Φ** relative to that of all species (function-function matrix **Φ**), which we call multifunctional participation index. Similarly, the colour of the edges quantify the weight relative to that of all species along the connections. (d) Multifunctional species ranking based on enriched metadata of the conditioned function-function networks. In order to rank species, we use the participation indices to obtain its keystone vector (see the text and Methods), which is shown in the figure, colour-coded to distinguish contributions from different functions.

First, we can explore how functions are intertwined at different resolution scales in such plant embedding by sequentially pruning edges in **Φ** and monitoring the size of the largest connected component (LCC): e.g. if pruning a low-weight link had the immediate effect of separating the network into two clusters of similar size, this would mean functions belonging to each cluster would play similar (intra-cluster) ecological roles, and dissimilar roles for each cluster. Conversely, if pruning low-weight links did not have a noticeable effect on the size of the LCC, this would mean that all functions have similar ecological roles. Now, edge pruning can be performed in several ways, where an obvious choice is to sequentially remove edges in ascending link weight, allowing us to evaluate the role of so-called *weak ties*^38^ as potential assemblers. Conversely, one can also remove edges with higher weight first (see Methods). In **Fig. 4b** we plot how the (normalised) size of the LCC, S(m)/n, of the pruned **Φ** as pruning is performed, for both pruning strategies. Results show that at first order the function-function network is “robust” against plant species extinctions –as it remains mostly unaltered until over 50% of the edges have been pruned-, i.e. functions perform similar roles in the plant-mediated similarity space. Interestingly, we also find that edges with lower weight – pairs of functions which are more dissimilar in this plant-mediated projection, i.e. weak ties– tend to be slightly more critical for the network assembly, as the net difference in Area Under the Curve (AUC) between both pruning strategies is about +9%. This result seems in agreement with the theory of weak ties^38^ –originally put forward in the context of social systems, and studied theoretically in models of correlated networks^39^-, where less homophilic ties e.g. tend to be more relevant for network assembly.

In order to further inspect the clustered structure of functions in this plant-feature embedding, we also performed a hierarchical clustering of the system, by interpreting the inverse 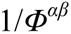 as the entries of a dissimilarity matrix. Results (**EDF2**) show that functions cluster together forming an intertwined core, with seed dispersal (and to some extent, fungal pathogenicity) being more peripheral.

Second, we already noted that each term 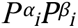 in the summation of 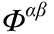 encodes the contribution of each specific plant species *i* to the connection (similarity) between functions α and β. Thus, the specific role of species *i* as a broker of functions can subsequently be disentangled from **Φ** by conditioning **Φ** only on a particular plant species *i*, and then appropriately visualised as a weighted (hexagonal-shaped) 6-node network 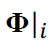, where (i) each node α represents a different function and is enriched with a node weight 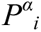 (according to the likelihood that the plant species *i* participates in that function), (ii) links between pairs of functions α and β denote that plant species *i* participates in both functions and the link weight 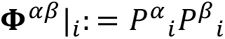 quantifies that contribution^b^: see **Fig. 4c** for some examples and **EDF3** for the complete set of networks {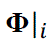}. The *metadata* (properly normalised sets of nodes and edges’ weights) of 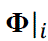 informs the (multifunctional) participation pattern of each plant species *i* in the whole ecosystem. Thus we use of such metadata to rank plant species’ multifunctional keystoneness. Since in this work we have a total of 6 observed functions, we associate to each plant species *i* a 6-dimensional vector (appropriately normalised), where each entry of the vector is 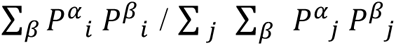: a *multifunction participation index* that quantifies the contribution to keystoneness of *i* via a particular function α (see Methods). The L_1_-norm of this vector thus provides the ranking ordering (see **Fig. 4d**). For better visualisation, in this figure we colour-code each element of the vector according to the function it corresponds to and plot the magnitude of each element as a coloured-segment of different length (the norm of the vector is thus the concatenation of line segments). This ranking summarises the role played by each plant species in assembling the ecosystem as propagated into the dual function-function representation (see also SI and **EDF4** and **EDF5** for complementary metrics). As we can see, some plant species such as *Withania frutescens* or *Lavatera maritima* play a critical role in every function and are thus at the top of the ranking (this agrees with complementary analysis in **EDF7** and **EDF9**). In contrast, some other species such as *Heliotropium sp.* or *Asparagus horridus* play a much weaker role.

Finally, it is important to clarify that the (multifunctional) species keystoneness index above does not correlate significantly with species relative abundance (Pearson r = 0.31, p-value = 0.24) or vegetation cover (Pearson r = 0.47, p-value = 0.06) (see **EDF6** and Extended Data Table **EDT1** for statistical details), suggesting that such keystoneness is neither directly related to potential sampling biases nor is a simple byproduct of species abundance, pointing to a more subtle property, unveiled here through our formalism.

### Dual Function-centric embedding: plant-plant networks and (multispecies) function keystoneness

The dual concept of *function keystoneness* can be explored following similar mathematical manipulations as in the phytocentric perspective: initially starting from MFEN, we project now onto the plant class (function-feature embedding) and thus construct a resource-resource (i.e. plant-plant) effective interaction network with *N=16* nodes and weighted adjacency matrix 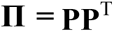, that leverages how *functions* broker the interaction between resources. Its elements 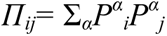 quantify the expected number of shared functions by two plant species, or alternatively how similar two plant species are in a function-feature embedding, see **Fig. 5a** where we again find that edge weights are heterogeneous. We then proceed to explore how functions act as plant species assemblers by performing a pruning analysis of **Π**. Results (**Fig. 5b**) suggest that plant species are more clustered: the pruning strategy of removing the edges with higher weight first (i.e. removing the links for the pairs of plants which show higher similarity in function-centric projection) leaves the plant-plant network unaltered until about 70% of the edges have been pruned, but conversely, the network smoothly dismantles after removal of 35% of the edges with lower weight. These findings reaffirm the significance of *weak ties* in this ecological context. In other words, plant-plant interactions that on average have a lower weight (i.e. are overall mediated by fewer functions or are observed less frequently or with different patterns, i.e. more dissimilar plant species) have a more critical role as plant-plant assemblers. Moreover, the net difference of AUC between both pruning strategies is about +16%, almost twice as large as that found for the function-function network, overall suggesting that functions have a more hierarchical role than plants. Finally, AUC for the plant-plant network descending-order pruning analysis (0.93) is greater than the respective AUC (0.84) for the function–function network. This implies that comparatively speaking, the plant-plant network is more cohesive (i.e. displaying more function-based similarity) that the other way around, i.e. functions assemble plants slightly more than the other way around.

**Figure 5:**
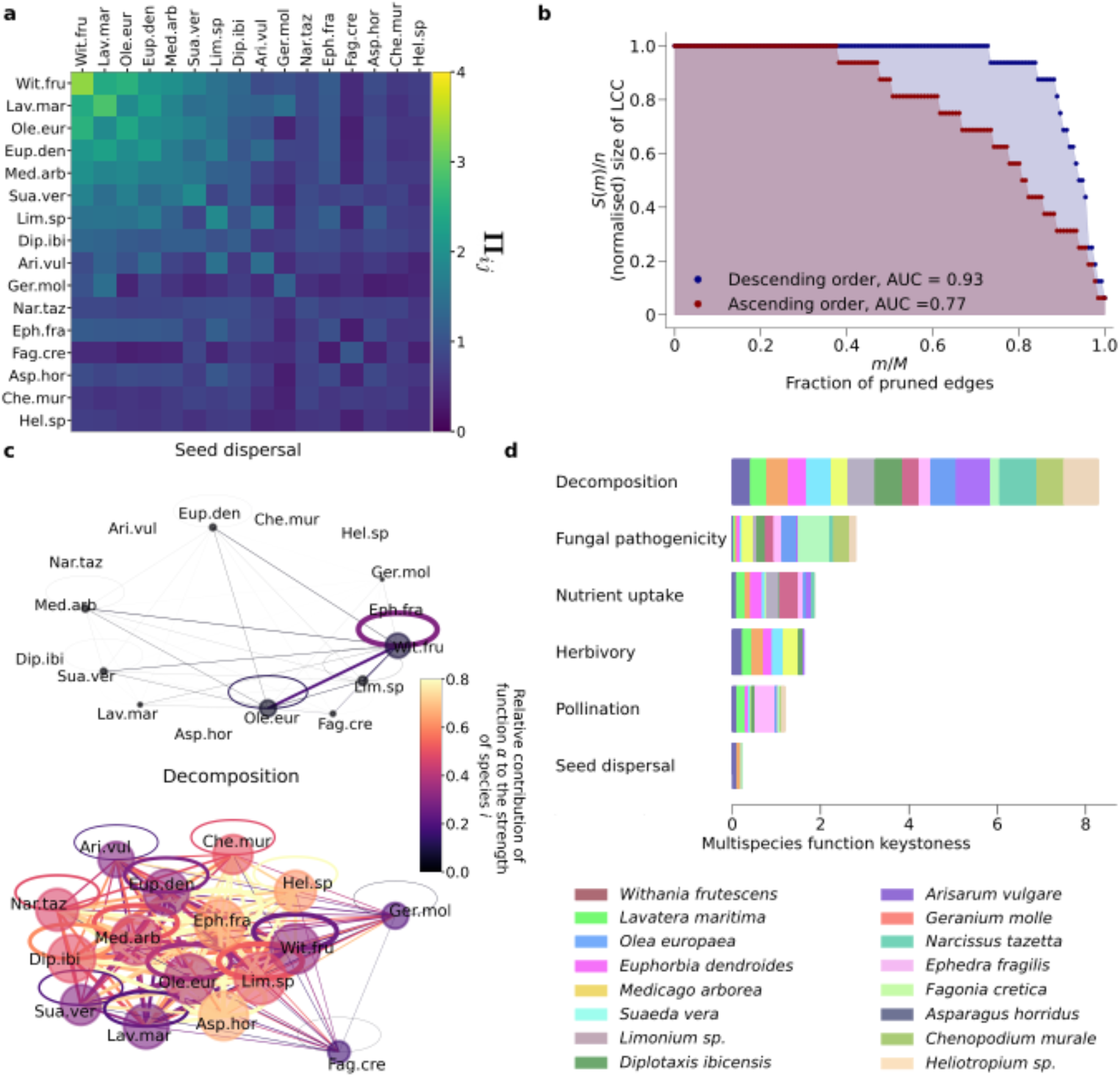
Function-centric embedding: plant-plant networks and multispecies function keystoneness. (a) Plant-plant network’s weighted adjacency matrix **Π** for the Na Redona dataset. (b) Pruning analysis of **Π,** performed as in Fig. 4b, finding qualitatively similar results as for the equivalent analysis on **Φ** although with a more emphatic role of weak ties. (c) Two examples of conditioned plant-plant networks **Π**|^*i*^ computed by conditioning on the functions decomposition (top) and seed-dispersal (bottom), showing two ecological functions with very different roles as species assemblers. (d) Multispecies function ranking based on enriched metadata of the conditioned plant–plant networks (see text and Methods). In the panel, for each function we plot its keystoneness vector (see Methods), colour-coded to distinguish contributions from different species.

We also analysed how plants cluster together at different resolution scales in this function-feature embedding by computing a hierarchical clustering on a dissimilarity matrix whose entries are 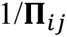, see **EDF7**. Interestingly, clusters of plants to reflect the multifunctional species keystonness found in **Fig 4.d**.

Finally, to evaluate and rank the role of each specific function as an ecosystem assembler, we replicate the analysis performed before and proceed to disaggregate **Π** by conditioning on each function, in order to extract 6 different plant-plant networks (one per function α) with weighted adjacency matrices 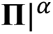 with elements 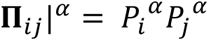 (see **Fig. 5c** for some examples^c^ and **EDF8** for the complete set 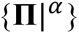). The metadata of 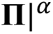 provides the plant participation of each function α and a ranking of *(multispecies) function keystoneness* can be derived (**Fig. 5d**). This ranking certifies the heterogeneity of roles and impacts of the different functions (to be compared with **EDF2**).

## Discussion

Our proposed RCF/MFEN framework offers a comprehensive and mathematically sound approach to unveil the complex architecture of multifunctional ecosystems. Let us first discuss and interpret in some detail the findings of such analysis as applied to the Na Redona dataset. Results of the phytocentric perspective indicate that plant species contribute hierarchically to multiple functions, i.e. they multitask^43^. Interestingly, the first six species in the keystoneness hierarchy, i.e. those with the highest multitasking indices, are all woody shrubs. The rest, except for *Ephedra fragilis*, are all herbaceous. Herbs such as *C. murale* and *Heliotropium* sp. play a minor role in linking the overall network, which does not mean they are not important for particular ecological functions. The finding that woody shrubs are those more strongly involved in different functions might be attributed to the longer lifespan of such species compared to herbs, which allows them to link to a wider array of species in each type of interaction. However, more in-depth studies are needed to unveil the exact mechanism of such multitasking. Interactions between plants and fungi (especially saprotrophic and pathogenic fungi) were found to play the most important role in assembling the multitrophic network. Microbial decomposers, together with plants and herbivorous insects, are also important drivers of ecosystem functioning in grasslands, where a positive association has been documented between richness or abundance and multiple ecosystem services^44^.

Results on the function-centric perspective suggest that saprophytic interactions (decomposition) exhibit a disproportionate prevalence: this agrees with a relatively recent shift in the interest and relevance of underground –in contrast to above-ground–ecological interactions. In any case, the pattern remains hierarchical (some plants dominate in one function while others play a dominant role in another or several functions), as quantified by our measures of multispecies function keystoneness. The finding that weaker function-function and plant-plant effective interactions (lower similarity ones) tend to play an important role in assembling the multifunctional ecosystem in these embeddings is reminiscent of the behaviours reported in social networks^38, 46^, where less frequently used interactions or less homophilic ones are those which tend to be information brokers, and the overall system tends to be more fragile against perturbations of these weaker links^39^.

Finally, it is worth noting that our whole framework is in principle metric-agnostic, i.e. while we used some specific quantification metrics, others alternatives are also possible^45^ (see SI for complementary metrics).

Now, it is important to acknowledge that there are certain limitations to our analysis and methodology, mainly associated with the unavoidable specificities of data collected in the Na Redona islet. First, the particular contraction method we used of the original tensor is based on the fact that in each layer, the interactions are bipartite; that is, there are no direct interactions between plants or between animals/fungi. Consequently, incorporating competition between plants and/or animals/fungi might eventually require some refinement of the framework. Second, while multifunctional, our collected data is eminently phytocentric, instead of zoocentric. If newer data allows us to account for purely zoocentric observations (ideally mixed plant-zoo-centric observations), retaining the complete RCF tensor would be convenient instead of contracting it. Third, since field observations were focused primarily on plants, we lack specific information, such as the trajectory of insect pollinators. Consequently, we cannot directly quantify the probability that a plant is included in the diet of a particular insect pollinator species. In this case, the parsimonious approach is to assume independence, and this is the base of our derivations. However, the contribution of a plant species to a function is typically mediated by the participation of several animal/fungal species, and the existence of correlations in the behaviour of different species is possible. Take for example, again, pollination. In our calculations we assumed that a universal probability governs whether a specific plant is visited or not. However, we do know that plants may attract specific pollinator species (especially the most specialised ones that have particular flower traits, e.g. long corolla tubes), resulting in a latent correlation kernel. Accordingly, the advent of new data should be used to test the validity of our parsimonious assumption.

We shall now provide some final remarks on our proposed framework. The RCF tensor serves as a powerful tool for capturing and quantifying the multidimensional nature of ecosystems. It facilitates data standardisation by integrating the multifunctional dimension into a resource-consumer*(–function)* paradigm and its contraction into a MFEN leads to an effective resource-function mapping, providing valuable insights into the interplay between the phytocentric and function-centric perspectives. Indeed, this framework is conceptually general and can be easily extended to characterise such interplay in other complex systems. For instance, it can be applied to genetics, where our resource–function mapping resembles a classical genotype-phenotype map^48^, illustrating how genes interact to give rise to phenotypes in various animals. Similarly, in economic systems, this interplay can shed light on how goods are traded among countries across different economic sectors^49^. The MFEN (and its projections) unveils the duality of plant species and interaction functions (phyto vs function-centric perspectives), where the concept of multifunctional species keystoneness and the dual concept of (multispecies) function keystoneness naturally emerge as two (interconnected) sides of the same coin. The former refers to species that play key roles to maintain a multifunctional system –with impact on e.g. network stability^43^-. In contrast, the latter refers to the different roles functions play in keeping species coexistence in an ecosystem. Both concepts are indeed intertwined, feeding back into each other. For instance, in the context of climate change, it is not uncommon to witness drastic and sustained temperature increases over large geographic regions. If such exogenous perturbations induce, e.g. phenological mismatches between flowering plants and insect pollinator visits, then the complete pollination function might be threatened before any of the species are themselves threatened^50^, i.e. perturbation takes place initially in the function dimension. It can then propagate with potential cascading effects for both species and other functions, as described in the resource–function map. Conversely, the decline or extinction of a seed disperser (animal species dimension) may trigger declines or plant extinctions, which in turn cascade to affect species involved in other ecological functions, such as underground fungi^51^. Furthermore, the extinction of a keystone plant species can also influence other species’ interactions, either by causing interaction rewiring or by modifying interaction strengths^52^. Such cascading effects across interaction types become particularly problematic in double-mutualistic interactions, i.e. when the same animal acts, for example, as pollinator and seed disperser of the same plant species^53^. Yet, not only the disruption of mutualistic interactions but also of antagonistic interactions could derive functional losses, e.g. the loss of top predators may indirectly affect ecosystem productivity and metabolism^54^. However, few studies still exist on how human activities can alter ecosystem multifunctionality, both directly on ecosystems and indirectly through the loss of multifunctional biodiversity^6, 55^. Moreover, how organisms at different trophic levels interact to influence ecosystem multifunctionality in the presence of multiple concurrent anthropogenic drivers remains largely unexplored^44, 56^. Thus, we hope our framework offers a promising approach for evaluating the relative vulnerability of ecosystem functions to anthropogenic stressors. As we delve deeper into understanding ecosystems, the integration of temporal, spatial, and dynamical features within this framework, coupled with its application to ecological data from other environments, emerge as exciting avenues for future research.

## Methods

### Study site and field sampling

The fieldwork was carried out on Na Redona (39^°^10^’^5 ^“^ N, 2^°^58^’^35^”^ E), an islet of approximately 11 ha and 56 m high in the Cabrera Archipelago National Park (Balearic Islands, Western Mediterranean Sea). Its primary habitat is Mediterranean shrubland with a relatively rich plant species diversity (*ca.* 108).

In two contrasting seasons, a team of five people visited the islet for five consecutive days at the peak of flowering (April/May) and at the peak of fruiting (October/November), to sample the different types of interactions between plants and (1) pollinators, (2) herbivores, (3) seed dispersers, and root-colonising (4) saprotrophic, (5) symbiotic, and (6) pathogenic fungi. In each sampling season, six transects (100 m long x 10 m wide and separated from each other by at least 100 m) were established to cover the main microhabitats and the entire altitudinal gradient of the island. In such transects, we assessed plant abundance (number of individuals) and vegetation cover (m^2^) of each plant species, and we recorded all species interactions except the seed-dispersal ones (see below).

Plant-fungi interactions were sampled by collecting the roots of five individuals of each plant species along the transects. Dry-cleaned roots were immediately preserved in silica gel until processing for DNA extraction. Then, PCR amplification was performed using fungal-targeted ITS1 primers ITS1f-ITS2^57, 58^. PCR products were cleaned, quantified, and sequenced at an equimolar concentration (Illumina sequencing using 2×350 bp MiSeq) at GENYO (University of Granada, Spain). Reads were filtered and processed using QIIME 2 pipeline^59^, and assigned to fungal ASV (amplicon sequence variants) using DADA2^60^. Taxonomy and functional groups were determined using the UNITE database^61^ and FUNGuild^62^.

To sample plant-pollinator interactions, we conducted censuses (of 10 min of duration, and at different times of the day) consisting of direct observations on each plant individual. Each species was observed for at least one hour during the sampling season, although the total time depended on flower availability (mean = 2 h, total sampling effort across species = 29 h). In each census, we recorded any animal (i.e. insects, lizards, or/and birds) that contacted the reproductive parts of the flowers. Insect species were photographed and identified using the reference collection available at IMEDEA. When unknown, they were captured and later identified by an entomologist, to the species level when possible or to the morphospecies level otherwise. Additionally, we captured 30 lizards using a noose on the tip, and 26 land birds using mist nets, being the latter not very abundant and difficult to catch since the islet is dominated by open vegetation or/and shrublands. We sampled the potential pollen carried by each lizard and bird by swabbing a cube (approx. 3mm^3^) of fuchsine-stained glycerine jelly on their snout and beak and peri-mandibular feathers, respectively. The gelatine cube was then placed on a microscope slide, melted, and covered with a slip. The entire slide area was inspected under a light microscope to identify and count all pollen grains to species level using a pollen reference collection available at IMEDEA. Evidence of flower visitation was considered if more than six pollen grains of a given plant species were detected in the sample. Each captured individual was considered a sampling unit.

Plant-herbivore interactions were evaluated in five individuals of each plant species, browsing the isolated branches and recording all arthropods found feeding on plant tissues. Unknown herbivorous species were captured and later identified by an entomologist, to the species level when possible or to the morphospecies level otherwise-e.

Finally, to sample plant-seed disperser interactions, we collected droppings and pellets of gulls (236), droppings of passerines (21), and lizards (375), and anthill material (4). We identified all taxa to plant species level or morphospecies under a stereomicroscope, by comparing the seeds with a reference seed collection also available at IMEDEA.

### Estimation of the weights in the RCF tensor

Taking advantage of the simplicity of a small island, we apply the developed framework by focusing on six layers of complexity corresponding to six ecological functions: pollination, herbivory, seed-dispersal, decomposition of organic matter, nutrient uptake (mycorrhizas) and fungal pathogenicity. Saprotrophic fungi are the primary agents of plant litter decomposition, thus, key regulators of nutrient cycling. Mycorrhizas play a key role in terrestrial ecosystems as they regulate nutrient and carbon cycles, influencing soil structure and ecosystem multifunctionality. Up to 80% of plant N and P is provided by mycorrhizal fungi and many plant species depend on these symbionts for growth and survival. Fungal pathogenicity is not a standard ecological function per se but can impact ecological functions and processes through its effects on population dynamics, biodiversity, trophic interactions, nutrient cycling, succession, and coevolution. Understanding the interactions between fungal pathogens and their is indeed important for comprehending and managing the ecological consequences of these diseases^68, 69^.

Our common species in six layers are plants (‘resources’), and we study their interactions with ‘consumers’: (1) pollinators, (2) herbivores, (3) seed dispersers, (4) saprophytic, (5) symbiotic and (6) pathogenic fungi. We assume a one-to-one map from interaction type to function, i.e. every time an interaction is recorded, the type of function it belongs to is also annotated –leaving no room for ambiguity-regardless of the inherent interaction strength (e.g. the time elapsed in the interaction), so the effective interaction strength is given through their frequency in the data. We acknowledge the evidence to support the crucial role of inherently-weak interactions in ecology, as they can act as both stabilizing factors and drivers of unstable dynamics^63, 64^. Our choice does not account for the variability in inherent interaction strength, and a general method is needed to explore the robustness of the mapping. This is highlighted in the literature on interaction strengths in food webs, which emphasises the need for clarity in terminology and definition, as well as exploring new ways to estimate biologically reasonable model coefficients from empirical data^65^.

The strength of an interaction between plant species *i,* and animal species *x* at function *α* (i.e. at any given ecological function), was calculated as:

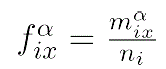

where 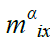 is the number of individuals of plant *i* on which an animal species *x* was detected along function α, and *n_i_* is the number of individuals of plant species *i* sampled. Plant–fungus interactions were quantified as the proportion of different taxa ASV reads per plant species^66^. For illustration, if say a total of *n_i_* =20 samples of the plant species *i* were monitored, and 12 of them were observed being pollinated by a single animal species whereas 10 of them were observed being dispersed (7 by animal species 1 and 3 by animal species 2), then the associated probabilities 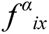 would be 12/20, 7/20, and 3/20 respectively (observe that these don’t add up to one: events are independent and thus probabilities are only normalised in terms of an event and its complementary). For example, in the case of the pollination layer, the weights of the interaction measure the fraction of plants of a given species that has been observed being pollinated, in other words, the probability that a random plant of species *i* is being pollinated.

### Resource-Consumer-Function tensor (RCF) visualisation

The specific visualisation of the RCF tensor presented in **Fig. 2**, containing plant species and the weight of interactions with animal/fungus species, was obtained using Netgraph, a freely available library implemented in Python (https://github.com/paulbrodersen/netgraph, last accessed: March 10, 2023). This layout also incorporates a study of the mesoscale organisation of the RCF while considering the network as a single-layer one. For that purpose, we used Infomap, a network clustering algorithm based on the Map Equation freely available at (https://github.com/mapequation/infomap.git, last accessed: March 10, 2023). The same clusters were also found using an alternative method (Louvain community detection algorithm). Generally, a plant species interacts with an animal/fungi species through a single ecological function. By projecting the tensor/multigraph onto a single layer, we focus the community detection algorithm’s attention on the plant-animal/fungi interactions. This allowed us to visualise 15 clusters, which broadly group the plant-animal/fungi pairs with the strongest interactions. We used different colours to identify each type of interaction and its associated animal/fungi species. For other visualisations, including Infomap enriched with multilayer properties, see **EDF2**.

Within our data, only 3 interactions between a plant species *i* and an animal/fungi *x* were catalogued as belonging to two different ecological functions. This happens, for example, when an insect exhibits herbivory during its larval stage and pollination during its adult stage. These interactions are represented as multilinks in the RCF, and are visualised with corresponding colours in the network without overlapping. Colours are assigned to the animal/fungi nodes based on one of the interactions, with a border of the colour corresponding to the other interaction.

### Estimation of edge weights P^α^_i_ in MFEN

To contract animals in the RCF tensor and build the MFEN, we follow a simple logical manipulation of probabilities. By construction, in the RCF the weight 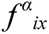 is the probability of finding a specific resource *i* and a specific animal/fungus *x* interacting via a specific function α, thus 1 –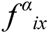 (the negation) is the probability of *not* finding [a specific resource i and a specific animal/fungus *x* interacting via a specific function α]. Assuming independence of events, the probability of not finding a specific resource *i* and *any* animal/fungus *x* interacting via a specific function α is, consequently, 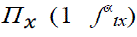. We finally negate this again, so the probability of finding a specific resource *i* and *any* animal/fungus *x* interacting via a specific function α is 1 – 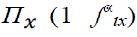. This probability is independent of *x*, it just describes the probability of a resource *i* effectively interacting via a function α, i.e. the probability that a plant species *i* participates in a function α. We call this the participation strength 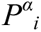.

### Generalist-Specialist ranking

The ranking of generalist and specialist species and functions in the bipartite network provides insight into the organisation and functioning of complex ecological systems. In particular, we evaluated the strength contribution of nodes belonging to the function class relative to the overall strength of all nodes in the class. The strength of a node is calculated by adding the weights of all its edges. For a given function α, the strength of the function is obtained as 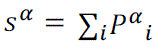. Then, the overall strength of the function class can be calculated by summing the strengths of all the individual functions, which can is expressed as 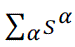.

Analogously, plant species’ class nodes are also ranked based on their strength contribution relative to the overall strength of the species in the bipartite network. The strength of a plant species *i* is obtained as 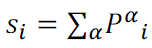. Then, the overall strength of a species class can be calculated by adding up the strengths of all individual species 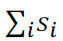. Note that to classify species as ecological function-generalists or function-specialists, both the number of functions in which they participate as well as the strength of their participation must be considered^37^.

### Φ and Π as similarity matrices

Let us consider **Φ** first. Mathematically, it is an 6 x 6 matrix as we have 6 different ecological functions displayed by the ecosystem. Each function-node α is enriched by a vector 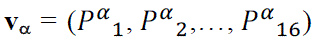, where **v_α_** accounts for the way in which function α interacts with each plant species. Accordingly, weights of the function-function network are given in terms of the scalar product 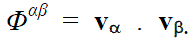 Since a scalar product can be interpreted as a similarity measure (correlation) between two vectors (with two ingredients: their magnitude and the angle between them), this simple mathematical observation indicates that 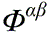 accounts for how similar two functions are, based on (i) how large or small is their participation strength across plants (vector magnitude), and (ii) how systematically mismatched (anticorrelated) is this pattern across plants (angle between vectors), i.e., it is function-to-function similarity measure emerging in a plant-feature embedding. The same can be said about **Π**: under this geometrical interpretation, *Π*_ij_ quantifies how similar plants *I* and *j* are, based on how similar their interaction patterns across functions are, i.e. it displays plant-to-plant similarities in a function-feature embedding.

### Pruning and robustness analysis

Starting from a weighted network, we order edge weights and proceed to prune the network in either ascending or descending order of edge weights. After pruning *m=1,2,…* edges, we compute the size (number of nodes) of the largest connected component of the network, obtaining a curve *S(m)*. *S(m)* is a monotonically decreasing curve, whose profile is indicative of the hierarchical structure of the network, and the area under this curve (AUC) determines the response of the network to the perturbation, where higher values of AUC mean that the network is more robust to a specific pruning strategy. To properly compare AUCs for different networks, AUC is computed by normalizing *S(m)* over the total number of nodes *N*, and m is normalised by dividing it over the total number of edges *M*. We compare ascending and descending pruning strategies by computing the enclosed area between the two curves, i.e. the difference between both AUCs, using a quadrature rule.

### Multifunctional participation index and associated keystoneness

The importance of resource *i* in connecting ecological functions is calculated with the multifunctional participation index (MPI)^67^. Let the strength of function α in resource *i* is 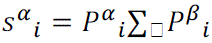 and the total connectivity of all plant species (all conditioned function-function networks) through the same ecological function be 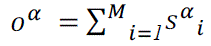. The MPI of plant species *i* and function α is obtained as 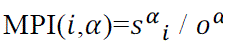. The resulting keystoneness vector of plant species *i* is composed of the 6 MPIs, one for each function (each of these MPI is colour-coded in **Fig. 4d**). The L_1_-norm of such vector (i.e. the sum of all entries) allows us to rank species accordingly (**Fig. 4d**). The same methodology is reproduced in the function-centric case by using the change of variable α→i. In this way, we can compute a multispecies participation index 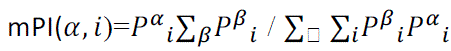. The resulting keystoneness vector of function α is composed of the 16 respective MPIs, one for each plant species (each of these mPI is colour-coded in **Fig. 5d**). The L1-norm of such vector (i.e. the sum of all entries) allows us to rank functions accordingly (**Fig. 5d**).

## Acknowledgements

We thank Xavier Canyelles that identified all insects; José Antonio Morillo that contributed to the bioinformatic analysis of plant-fungi interactions, Aarón González-Castro and Juan Pedro González-Varo for their help in the field, and Araceli Guillem Bosch for her help in the laboratory processing of pollen samples. M.C.B., L.L., and V.M.E. acknowledge funding from the Spanish Research Agency (MCIN/AEI/10.13039/501100011033) via project MISLAND (PID2020-114324GB-C22), the María de Maeztu project CEX2021-001164-M and project DYNDEEP (EUR2021-122007). S.H.P and A.T. acknowledge funding by MCIN/AEI/10.13039/501100011033 (CGL2017-88122-P, PID2020-114324GB-C21) and the María de Maeztu project CEX2021-001198-M. The study is also framed within project 101054177 *IslandLife* funded by ERC AdG to A.T. I.D is supported by a Marie Curie Postdoctoral Fellowship (HORIZON-TMA-MSCA-101068643). R.H. was funded by the Portuguese Foundation for Science and Technology (UID/BIA/04004/2020).

## Code availability statement

All the necessary codes will be available upon publication in a GitHub repository.

## Data availability statement

Data will be available upon publication in a GitHub repository.

## Author contributions

A.T., M.N., R.H., S.H-P. and S.R-E conceived and designed the research to build the multilayer network and gathered data in the field. S.H-P. led the data curation after all taxa were identified and built the multilayer network. S.R-E did the classification of fungi into the three functional groups. L.L. and V.M.E. conceptualised and lead the mathematical modelling. M.C-B. contributed to mathematical modelling, performed all the network analysis, data analysis and simulations and generated all the figures. S.H-P, M.C-B, L.L, A.T., I.D., C.M. and V.M.E discussed results. L.L., M.C-B and V.M.E. wrote the first draft, and all authors contributed to write and revise the final draft.

## Competing interest declaration

The authors declare no competing interests.

## Footnotes

a Note the slight abuse of notation in calling **P** a matrix instead of a rank-2 tensor: throughout this work, for simplicity, we shall call tensor only rank-3 tensors –as the RCF–, whereas all rank-2 tensors are called (adjacency) matrices, regardless the number of covariant and contravariant indices.

b This is compatible with the interpretation that the probability that a resource *i* is observed participating in two functions could be obtained from the product of the probabilities, as in Fig. 3a. Such no-correlation scenario leads the resulting matrix to be in the universality class of so-called rank-1 networks^40^, and can also be understood under the paradigm of fitness-mediated good-get-richer networks^41^ or a particular class of hidden variable network^42^.

c The conditioned plant-plant networks are again in the universality class of rank-1 networks40 and can be interpreted as fitness-based or hidden variable networks^41, 42^.

## Notes

### Competing Interest Statement

The authors have declared no competing interest.

